# Heptanol-mediated phase separation determines phase preference of molecules in live cell membranes

**DOI:** 10.1101/2021.10.18.464904

**Authors:** Anjali Gupta, Danqin Lu, Harikrushnan Balasubramanian, Zhang Chi, Thorsten Wohland

## Abstract

Plasma membranes contain diverse nanoscale assemblies of lipid and protein domains. Specific localization of lipids and proteins in these domains is often essential for membrane function and integrity. Due to the nanoscale size and dynamic nature of membrane domains, identification of molecules residing in domains either is not possible with modern imaging techniques or requires advanced methods with high spatiotemporal resolution. Such methods need expensive equipment making these approaches inaccessible and thus difficult to implement at large scale. Here, we present a novel membrane fluidizer-induced clustering (MFIC) approach to identify the phase-preference of molecules in intact cell membranes. Experiments in phase-separated bilayers and live cells on molecules with known phase preference demonstrate that heptanol hyperfluidizes the membrane and stabilizes phase separation in cell membranes. The domain stabilization results in a transition of nano-to micron-sized clusters of associated molecules and allows identification of molecules localized in domains using routine microscopy techniques. This assay can be carried out on both genetically and extrinsically labelled molecules in live cell membranes, does not require any invasive sample preparation and can be carried out in 10-15 minutes. This inexpensive and easy to implement assay can be conducted at large-scale and will allow easy identification of molecules partitioning into domains.

## INTRODUCTION

Plasma membranes (PMs) are complex entities composed of a diversity of lipids and proteins that associate in different combinations, thereby resulting in plasma membrane heterogeneities. These heterogeneities exist in the form of protein clusters, lipid-lipid complexes, or combinations thereof, e.g., cholesterol sphingolipids or protein-lipid complexes^1,2^. In addition to the preferential interaction of these molecules with each other, there is a dynamic cytoskeleton network and extracellular matrix (ECM) that influences the spatial organization of heterogeneities in the PMs^3,4^. PMs are extremely dynamic and are highly susceptible to change their physical properties and organization. Upon interaction with membrane-active compounds, such as peptides or anesthetics, membranes undergo a reversible modulation of spatial organization and membrane order^5–9^. Membrane proteins often reside in a specific lipid environment in their resting state, and they change their environment upon activation^10^. With increasing evidence, it is now realized that dynamic changes in the lipid environment of these proteins are essential for their functionality and regulation. The lipid environment of signalling proteins is hypothesized to influence the signal transduction originating at the plasma membrane^11–13^. It was recently shown that specific structural characteristics of proteins such as palmitoylation, length of the transmembrane segment, and type of amino acids in the transmembrane region could determine their preference for a certain phase^14^. However, the identity of the lipid environment surrounding the signalling proteins remains controversial, primarily because most of the existing literature relies on data obtained from indirect and artifact-prone methods^15,16^. The reason why indirect methods have been used to detect membrane domains is that the size of domains is typically below the diffraction limit, and thus, they are inaccessible by routine imaging methods. Moreover, due to their dynamic nature, membrane domains typically last from microseconds to seconds and are often difficult to detect. To examine the complex plasma membrane structure and dynamics, artificially reconstituted model membranes have contributed significantly^17–26^. However, they cannot recapitulate every physiologically relevant attribute. For instance, although model membranes can be tuned to exhibit micron-sized domains of a specific phase (liquid disordered and liquid-ordered), it is nearly impossible to visualize domains directly in cell membranes as the domains in plasma membranes are much smaller and are very sensitive, e.g., even giant plasma membrane vesicles (GPMVs) do not keep the same organization as a live cell plasma membrane^1,14,27^. Therefore, there is a clear need of simple methods that can detect the phase preference of membrane proteins in live intact cell membranes. Biochemical methods that have been used to differentiate the proteins that localize in the raft and non-raft phases include detergent-resistant extraction^28^, immunostaining^29^, cell fractionation followed by mass spectrometry^30^. However, these methods are artifact-prone as either they involve the use of non-physiological experimental conditions or require fixed samples. Due to these reasons, fluorescence-based methods combined with live-cell imaging and spectroscopy (e.g. FCS diffusion laws) are alternatives for determining the membrane heterogeneities^31–35^. In previous studies, phase-specific fluorescent dyes and protein anchors have been used to understand the dynamic properties of the individual phases in live-cell membranes^33,36^. Despite the wide usage of such methods they have been difficult to implement due to the requirement of specialized instrumentation and they also have some exceptions that pose problems in their interpretation^37–39^.

In this work, we present a novel MFIC methodology to determine the localization of molecules in live cell membranes. In this study we use heptanol as a membrane-fluidizing agent and show using total internal reflection fluorescence microscopy (TIRFM) that there is reversible reorganization in the cell membrane as a result of heptanol treatment. The molecules that reside in cholesterol-dependent domains segregate into micron-sized clusters, while molecules that reside outside the cholesterol-dependent domains stay dispersed. We test this assay in both model membranes and live intact cell membranes using several molecules with known phase preference. Moreover, we use this method to probe the localization of signalling related proteins such as epidermal growth receptor factor (EGFR), IL-2Rα, KRas, and H-Ras. Furthermore, we test this method on other cell lines to ensure the universality of this method across different cell membranes. This method is an inexpensive, fast (~ min), and minimally invasive way to determine whether a molecule resides within lipid domains in live cells.

## EXPERIMENTAL PROCEDURES

### Reagents

The lipids 1,2-dioleoyl-sn-glycero-3-phosphocholine (DOPC), 1,2-dipalmitoyl-sn-glycero-3-phosphocholine (DPPC) and cholesterol (Chol) were used in this work. Head group labelled rhodamine dye 1,2-dimyristoyl-sn-glycero-3-phosphoethanolamine-N-(lissamine rhodamine B sulfonyl) (ammonium salt) (RhoPE; 14:0 Liss Rhod PE) was used as the fluorophore to label supported lipid bilayers. The lipids and dye were purchased from Avanti Polar Lipids (Alabama, USA) and dissolved in chloroform.

1,1’-dioctadecyl-3,3,3’,3’-tetramethylindocarbocyanine perchlorate (DiI-C_18_, #D3911), octadecyl rhodamine B chloride (R18, #O246) and CTxB-555 (cholera toxin subunit B (Recombinant) with Alexa Fluor 555 conjugate, #C34776) were purchased from Invitrogen (Thermo Fisher Scientific, Singapore). They were dissolved in anhydrous dimethyl sulfoxide (DMSO; #276855, Sigma-Aldrich, Singapore) to prepare the stock solutions. The fluidizer 1-Heptanol 98% (#H2805) was purchased from Sigma-Aldrich (Singapore).

An Alexa Fluor 488 conjugated EGFR monoclonal antibody was purchased from Cell Signaling Technology (EGF Receptor (D38B1) XP Rabbit mAb (Alexa Fluor 488 Conjugate), #5616S, Massachusetts, USA).

### Supported lipid bilayer (SLB) preparation

All glassware (slides, coplin jars, and round-bottom flasks) were cleaned thoroughly with an alkaline cleaning solution (Hellmanex III, Hellma Analytics, Müllheim, Germany) using sonication (Elmasonic S30H, Elma Schmidbauer GmbH, Singen, Germany) for 30 minutes. They were then washed with ultrapure water (Milli-Q, Merck, New Jersey, United States), submerged in 2M sulphuric acid, and sonicated again for 30 minutes. After washing the glassware with deionized water and immersing them in the water, a final sonication was done for another 30 minutes.

A silicone elastomer (SYLGARD 184 Silicone Elastomer Kit, Dow, Michigan, USA) was filled in an O-ring mould and cured at 65°C overnight. The O-rings were carefully removed using forceps and attached to a slide using the silicone elastomer, followed by curing at 65°C for 3 hours.

500 μM DOPC:DPPC:Chol (4:3:3) solution and 100 nM RhoPE were mixed thoroughly in a round-bottom flask and evaporated in a rotary evaporator (Rotavapor R-210, Büchi, Flawil, Switzerland) for 3-4 hours until a thin lipid film was formed. The lipid film was dissolved in 2 ml of a buffer solution (10 mM HEPES, 150 mM NaCl, pH 7.4) and sonicated until the solution became clear, indicating the formation of large unilamellar vesicles. 200 μl of the lipid solution was added into an O-ring attached to a slide and incubated at 65°C for 1 hour to allow vesicle fusion and formation of the SLB. The SLB was cooled to room temperature (25°C) for 30 minutes and then washed with the buffer solution multiple times to eliminate the unfused vesicles. SLB measurements were done at 37 °C.

### Plasmids

The green fluorescent protein-tagged glycosylphosphatidylinositol-anchored protein (GPI-GFP plasmid) was a kind gift of John Dangerfield (Anovasia Pte Ltd, Singapore). The plasmids IL2Rα-EGFP (Addgene plasmid #86055), mEGFP-HRas (Addgene plasmid #18662), pLVET-HA-K-RasG12V-IRES-GFP (Addgene plasmid #107140) were purchased from Addgene (Massachusetts, USA). The construction of EGFR-mApple has been previously described^40^. The EGFR-mEGFP plasmid was constructed in the same way as EGFR-mApple.

The sequences of the transmembrane domain of linker for activation of T-cells (WT-trLAT) and a mutant with all the transmembrane domain amino acids (except the palmitoylation sites) mutated to leucines (allL-trLAT) have been previously described^41^. DNA duplexes were designed with the trLAT sequences flanked by AgeI and SpeI restriction sites on the 5’ and 3’ ends, respectively, with a 6-base linker between them. The duplex DNA sequences were synthesized and purchased from Integrated DNA Technologies Pte. Ltd. (Singapore). The EGFR-mApple and EGFR-mEGFP plasmids were digested with AgeI (AgeI-HF, R3552S, New England BioLabs, Massachusetts, USA) and SpeI (SpeI-HF, R3133S, New England BioLabs, Massachusetts, USA) to create the plasmid backbones. The WT-trlAT and all-trLAT sequences were also digested with AgeI and SpeI to create the inserts. The backbones and inserts were ligated using T4 DNA ligase (M0202S; New England BioLabs, Massachusetts, USA) to create four plasmids – WT-trLAT-mEGFP, WT-trLAT-mApple, allL-trLAT-mEGFP, and allL-trLAT-mApple.

### Cell culture

The protocol detailing the steps in the preparation of live-cell samples for fluorescence applications is provided in Protocol Exchange^42^. SH-SY5Y (#CRL-2266) and HeLa (#CCL-2) cells were obtained from ATCC (Manassas, Virginia, USA). They were cultivated in Dulbecco’s Modified Eagle Medium (DMEM/High glucose with L-glutamine, without sodium pyruvate – #SH30022.FS; HyClone, GE Healthcare Life Sciences, Utah, USA) supplemented with 10% fetal bovine serum (FBS; #10270106, Gibco, Thermo Fisher Scientific, Singapore) and 1% penicillin-streptomycin (#15070063, Gibco, Thermo Fisher Scientific, Singapore), at 37 °C in a 5% (v/v) CO_2_ humidified environment (Forma Steri-Cycle CO_2_ incubator, Thermo Fisher Scientific, Singapore).

Cell cultures that were ~90% confluent were passaged. The spent media was removed from the culture flask and 5 ml 1X PBS (phosphate-buffered saline; without Ca^2+^ and Mg^2+^) was used to wash the cells twice. 2 ml TrypLE Express Enzyme (1X; #12604021, Gibco, Thermo Fisher Scientific, Singapore) was added, and the flask was placed in the CO_2_ incubator for 2-3 minutes. Upon detachment of the cells, 5 ml culture media was added to the flask to inhibit trypsin. The cell suspension was centrifuged (#5810, Eppendorf, Hamburg, Germany) at 200 × g for 3 minutes. The supernatant was discarded, and the cell pellet was resuspended in 5 ml 1X PBS. An automated cell counter (TC20, Bio-Rad, Singapore) was used to count the cells, and the required number of cells was used for the next step of cell membrane staining or transfection.

### Cell membrane staining

After passaging, the required number of cells were seeded onto culture dishes (#P35G-1.5-20-C, MatTek, Massachusetts, USA) containing culture media and allowed to attach for 24 hours. DiI-C_18_, R18, and CTxB-555 stock solutions were diluted to 100 nM working concentration in imaging medium (DMEM with no phenol red, #21063029, Gibco, Thermo Fisher Scientific, Singapore) supplemented with 10% FBS. They were used for cell membrane staining. The media was removed and replaced with the working dye solution. The cells were placed in the CO_2_ incubator for 20 minutes. Then the dye solution was removed, and the cells were washed with 1X HBSS (Hank’s Balanced Salt Solution, with Ca^2+^ and Mg^2+^; #14025134; Gibco, Thermo Fisher, Singapore) twice. DMEM without phenol red (#21063029; Gibco, Thermo Fisher Scientific, Massachusetts, USA), hereafter called imaging media, was supplemented with 10% FBS and added to the cells before measurements.

### Transfection

After passaging, the required number of cells was centrifuged at 200 × g for 3 minutes. The supernatant was discarded, and the cells were resuspended in R buffer (Neon Transfection Kit, Thermo Fisher Scientific, Singapore). Suitable amounts of plasmids were mixed with the cells for transfection. The cells were electroporated according to the manufacturer’s protocol (electroporation settings: SH-SY5Y – pulse voltage = 1,200 V, pulse width = 20 ms, pulses = 3; HeLa – pulse voltage = 1,005 V, pulse width = 35 ms, pulses = 2) using Neon Transfection system (Thermo Fisher Scientific, Singapore). After transfection, the cells were seeded onto culture dishes containing DMEM (with 10% FBS; no antibiotics). The cells were incubated in the CO_2_ incubator for 20-48 hours before the measurements.

### Cell measurements

The transfected cells were washed twice with HBSS and imaging media (with 10% FBS) was added before measurements. EGFR-transfected cells were starved for a few hours in imaging media (without FBS to prevent aberrant activation of EGFR) before measurements. Stock heptanol solution was filtered with a 0.2 μm syringe filter and added to the imaging media to obtain the working concentration of 5 mM. Measurements were done after 10-20 minutes of incubation.

For the two-colour EGFR antibody measurements, the antibody was diluted 1:200 in imaging media and added to EGFR-transfected cells. The cells were incubated for 3 hours in the CO_2_ incubator. Subsequently they were washed with HBSS twice and imaging media was added. After initial imaging of the antibody labelling, 5 mM heptanol was added to the cells and they were imaged.

### Instrumentation

An inverted epi-fluorescence microscope (IX83, Olympus, Singapore) with a motorized TIRF illumination combiner (cellTIRF-4Line IX3-MITICO, Olympus, Singapore) and an oil-immersion objective (100×, NA 1.49, Apo N, Olympus, Singapore) was used for the imaging measurements. 488 nm (LAS/488/100, Olympus, Singapore) and 561 nm (LAS/561/100, Olympus, Singapore) lasers were connected to the TIRF illumination combiner. The laser power (as measured at the back aperture of the objective) used for both the 488 nm laser and the 561 nm laser was ~0.3 mW for the single-channel imaging and ~0.1 mW for two-colour imaging. The fluorescence emission was passed through a dichroic (ZT 405/488/561/640rpc, Chroma Technology Corp, Vermont, USA) and emission filter (ZET405/488/561/640m, Chroma Technology Corp, Vermont, USA) to an electron multiplying charge-coupled device (EMCCD; iXon^EM^+ 860, 24 μm pixel size, 128×128 pixels, Andor, Oxford Instruments, UK) camera. For the dual-colour measurements of antibody labelling of EGFR a dual-emission image splitter (OptoSplit II, Cairn Research, Faversham, UK) equipped with an emission dichroic (FF560-FDi01, Semrock, New York, USA) and band-pass emission filters (FF03-525/50-25 and BLP01-568R-25, Semrock, New York, USA) was used.

The ITIR-FCS measurements were carried out using another TIRF microscope (IX-71, Olympus, Singapore) equipped with an oil-immersion objective (PlanApo, 100×, NA 1.45, Olympus, Singapore) and an electron multiplying charge-coupled device (EMCCD; iXon 860, 24 μm pixel size, 128×128 pixels, Andor, Oxford Instruments, UK) camera. 488 nm (Spectra-Physics Lasers, California, USA) and 532 nm (Cobolt Samba, Sweden) lasers were used as the excitation sources that focused on the samples by a combination of two tilting mirrors and a dichroic mirror – 495LP (Omega Optical, Vermont, USA) for 488 nm and Z488/532M (Semrock, New York, USA) for 561 nm excitation, respectively. ~0.3 mW laser power was used for both lasers.

For the cell measurements, 37°C temperature and 5% CO_2_ atmosphere were maintained using an on-stage incubator (Chamlide TC, Live Cell Instrument, South Korea). Andor Solis (version 4.31.30037.0-64-bit) was used for image acquisition. The following camera settings were used: mode of image acquisition = kinetic, baseline clamp = “on” to minimize the baseline fluctuation, pixel readout speed = 10 MHz, maximum analog-to-digital gain = 4.7, vertical shift speed = 0.45 μs, EM gain = 300. A region of interest (ROI) of size 5.04 × 5.04 μm^2^ containing 21 × 21 pixels was selected on a cell. For FCS measurements, a stack of 50,000 frames was collected by the EMCCD with 1 ms recording time for DiI-C_18_ and 2 ms for the other samples. For single-colour and dual-colour imaging, a stack of 10 frames was collected at 20 ms and 100 ms exposure time, respectively.

### Data analysis

#### Estimation of diffusion coefficient from ITIR-FCS

After obtaining the image stacks, data analysis was performed using a home-written GPU-accelerated Imaging FCS 1.52 plugin in ImageJ^40,43,44^. A step-by-step protocol detailing FCS data analysis is provided in Protocol Exchange^43^. The fluorescence fluctuations at all pixels were calculated with the autocorrelation functions (ACFs) and fitted with the following equation after an exponential of polynomial bleach correction. Finally, quantitative maps of diffusion coefficient (D) were obtained as well as the FCS diffusion law intercept (τ0) performed by the FCS diffusion law analysis of the same image stack.

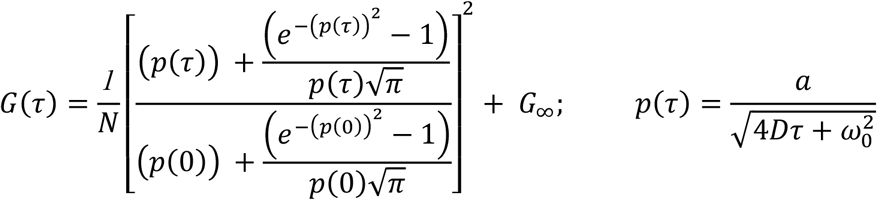

Here G(τ) is the theoretical model of the ACF in dependence of the correlation time (τ). *a* is the pixel side length, ω0 is the 1/e^2^ radius of the Gaussian approximation of microscope point spread function, while N is the number of particles. N, D and G_∞_ were all fitting parameters.

#### Cluster area fraction calculation

For calculation of the fractional cell area occupied by protein clusters (Figure S1), intensity thresholding was applied using Fiji. The first thresholding step was to demarcate the cell from the background using a simple intensity threshold (Image ➔ Adjust ➔ Threshold). The second step was to threshold the clusters using the Renyi entropy algorithm^45^ (Image ➔ Adjust ➔ Auto Threshold ➔ RenyiEntropy). The algorithm had trouble choosing a threshold to identify clusters when the intensity differences between cluster and non-cluster areas were small. To mitigate this, cells with high intensity differences between the cluster and non-cluster areas were processed as 16-bit images while cells with small intensity differences were converted and processed as 8-bit images.

The cells in resting state showed negligible clustering and the algorithm failed to pick up any clusters. Therefore, for these cells, the thresholding was done manually to select the few clusters that were present.

The fractional area occupied by the clusters on the cell was calculated as:

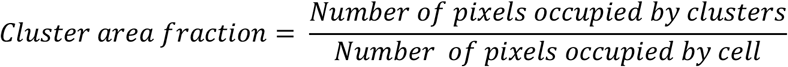

#### Cluster size calculation

For calculation of the average cluster size (Figure S1), all the images were converted to 8-bit and intensity thresholding was done using the Renyi entropy algorithm (except for resting cells which were manually thresholded) as explained in the previous section. While the 8-bit conversion eliminated tiny dim clusters and biased towards choosing the larger and brighter clusters, it allowed better separation of closely-spaced clusters resulting in more accurate average cluster size estimation. After thresholding, the clusters were analyzed in Fiji (Analyze ➔ Analyze Particles) to determine the average cluster size. The results from Fiji had units of squared pixel which was reported in Figure S1 after conversion to nm by calculating the square root and multiplying by pixel size (240 nm).

#### Statistical analyses

For the results in Figure S1, two-tailed homoscedastic T-test with 95% confidence interval was performed in Microsoft Excel. One-way ANOVA analysis was performed using an online calculator (https://www.statskingdom.com/180Anova1way.html). Tukey HSD post-hoc test was also performed with 95% confidence interval to identify the difference levels between group pairs in ANOVA.

## RESULTS AND DISCUSSION

### Heptanol induces hyperfluidization and domain segregation in model and live cell membranes

Fluidizers increase the membrane order and alter the organization of membrane components. Previous work demonstrated that benzyl alcohol (BA), a routinely used anesthetic, hyperfluidizes the membrane and induces the reorganization of domains^46^. A recent work showed that propofol also reorganizes cholesterol-dependent domains in cell membranes^47^.

We, therefore, investigated the effect of heptanol, an alcohol-based anesthetic, on membrane dynamics and organization. To systematically evaluate the impact of heptanol on the organization and fluidity of membranes, we tested its effect on a ternary lipid bilayer exhibiting phase properties closer to cell membranes and subsequently on intact cell membranes. We measure the fluidity of the membrane using imaging total internal reflection fluorescence correlation spectroscopy (ITIR-FCS), an imaging FCS modality that allows measurement of spatially-resolved molecular diffusion over a whole region of interest in a single measurement.

DOPC:DPPC:Chol (4:4:3) lipid bilayers exhibit coexisting liquid-ordered (L_o_) and liquid disordered (L_d_)^48^ phases, mimicking the phase behavior of an intact cell membrane. As observed in a resting cell membrane, the size of domains in this bilayer is below the diffraction limit, thus it is optically homogeneous (Figure 1a). To test the effect of heptanol on this bilayer, we treated the bilayer with 5 mM heptanol and performed ITIR-FCS measurements on them immediately after the treatment. Our results show that upon heptanol treatment, the D of Rho-PE increases from 1.21 ± 0.11 μm^2^/s to 3.51 ± 1.03 μm^2^/s (Figure 1a,b). Thus, as expected, heptanol fluidizes the membrane as inferred by an almost three times increase in *D*. Interestingly, we observed phase separation in the bilayer as there was formation of micron-sized lipid clusters 10 minutes after heptanol addition (Figure 1a,b).

**Figure 1:**
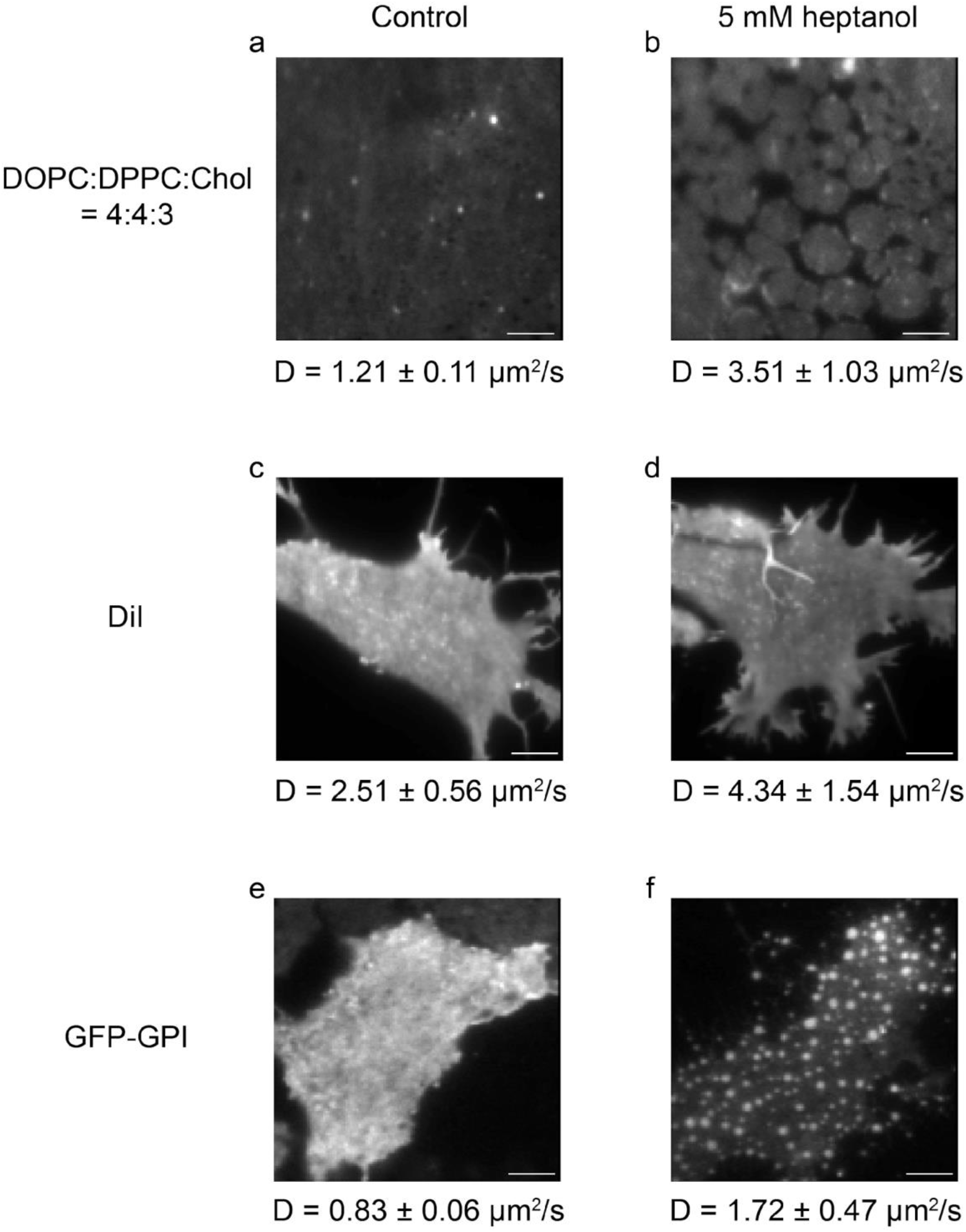
Heptanol induces hyperfluidization and domain segregation in model and live cell membranes. Samples were treated with 5 mM heptanol. Representative TIRF images and the corresponding average diffusion coefficients (D) of different samples are shown here. **(a,b)** Rhodamine-PE labelled DOPC:DPPC:Chol (4:4:3) supported lipid bilayer before and after heptanol treatment. The D increases from 1.21 ± 0.11 μm^2^/s to 3.51 ± 1.03 μm^2^/s following heptanol treatment. **(c,d)** DiI-C_18_ labelled SH-SY5Y cells before and after the heptanol treatment. The D increases from 2.51 ± 0.56 μm^2^/s to 4.34 ± 1.54 μm^2^/s following heptanol treatment. **(e,f)** GFP-GPI expressing SH-SY5Y cells before and after the heptanol treatment. The D increases from 0.83 ± 0.06 μm^2^/s to 1.72 ± 0.47 μm^2^/s following heptanol treatment. Experiments were repeated at least four times. The scale bars represent 5 μm.

Intrigued by this observation, we extended these experiments to live cell membranes. We tested the heptanol-induced organizational changes on molecular probes (DiI-C_18_ and GFP-GPI) with known phase preferences in cell membranes. Here we treat the cells with 5 mM heptanol and perform TIRF imaging and ITIR-FCS measurements. DiI-C_18_ localizes mainly in the fluid fraction of the membrane and has been used as a free diffusion marker, while GFP-GPI localizes in the cholesterol-dependent domains^10,24,33^. Heptanol treated SH-SY5Y cells labelled with DiI-C_18_ show an increase in *D* from 2.51 ± 0.56 μm^2^/s to 4.34 ± 1.54 μm^2^/s (Figure 1c,d). However, there was no visible aggregate formation in the cell membrane even 30 minutes post-treatment. In the case of GFP-GPI expressed in SH-SY5Y cells, heptanol treatment resulted in an increase in *D* from 0.83 ± 0.06 μm^2^/s to 1.72 ± 0.47 μm^2^/s (Figure 1e,f). After incubating the sample for 15 minutes, we observed phase separation on the cell membrane in the form of GPI clusters. Altered lateral segregation in the form of domain stabilization could be due to the immiscibility of domain components caused by hyperfluidity of the membrane.

### Heptanol stabilizes phase separation in live plasma membranes

Based on our results, we hypothesized that *heptanol-induced hyperfluidization stabilizes the phase separation in plasma membranes resulting in the formation of micron-sized clusters of the phases*, *as they exist at the nanoscale*. To test this hypothesis, we studied the effect of heptanol on the lateral segregation of molecules with known localization preference. Since heptanol induces microscopic changes in the lateral organization of molecules, TIRF images are sufficient to determine the molecular rearrangements on the membrane.

The B subunit of cholera toxin (CTxB) has emerged as a commonly studied domain-binding protein^24,49^. It binds to GM1-ganglioside, a ubiquitous cell surface glycolipid residing in the L_o_ phase of the membrane. To test the effect of heptanol on the lateral segregation of CTxB, we treated CTxB-labelled SH-SY5Y cells with heptanol and imaged them 15 minutes post-treatment. In this case, we observed the formation of clusters on the cell membranes (Figure 2a). CTxB forms non-specific clusters even in resting cell membranes. Thus, here we report the obvious increase in the number/size of clusters upon heptanol treatment. These experiments reveal the lateral segregation of endogenous GM1 molecules labelled by CTxB. Next, we probed the effect of heptanol on the organization of Rhodamine-C18 (R-18), a lipophilic dye containing a single 18-membered hydrocarbon chain. It is excluded from the L_o_ environments and thus is regarded as a marker of the L_d_ phase^36^. Owing to its lipophilic nature, this dye forms non-specific clusters in untreated cell membranes (Figure 2b). Upon heptanol treatment, we observed no additional cluster formation and the cell membrane remained homogeneous, even 30 minutes after treatment (Figure 2b). To further verify our hypothesis, we examined the effect of heptanol on the lateral segregation of two versions of the transmembrane domain (TMD) of the Linker for Activation of T-cells (trLAT). The partitioning properties of wild-type (WT) and various mutants of trLAT were extensively characterized by Lorent et al.^14^. It was shown that the WT-trLAT preferentially resides in the L_o_ phase of the cell membrane while the construct where all the TMD residues were mutated to leucine (trLAT-allL) showed dramatically decreased affinity to the L_o_ phase. Due to the existing knowledge regarding the partitioning of these constructs, we analyzed their organization upon heptanol treatment. We transfected these constructs in SH-SY5Y cells and recorded cell membrane images using TIRFM before and after treatment. In untreated cell membranes, WT-trLAT exhibits homogeneous expression while trLAT-allL is extremely sensitive to the total protein amount transfected in the cell and often shows clusters at higher protein concentrations (Figure 2c,d). The concentration of trLAT-allL plasmid was optimized to achieve a sufficient signal-to-noise ratio and minimal non-specific clustering. Consistent with our hypothesis, we observed that in the case of WT-trLAT, heptanol induced formation of clusters in the cell membrane while trLAT-allL did not undergo any additional clustering in response to heptanol treatment (Figure 2c,d).

**Figure 2:**
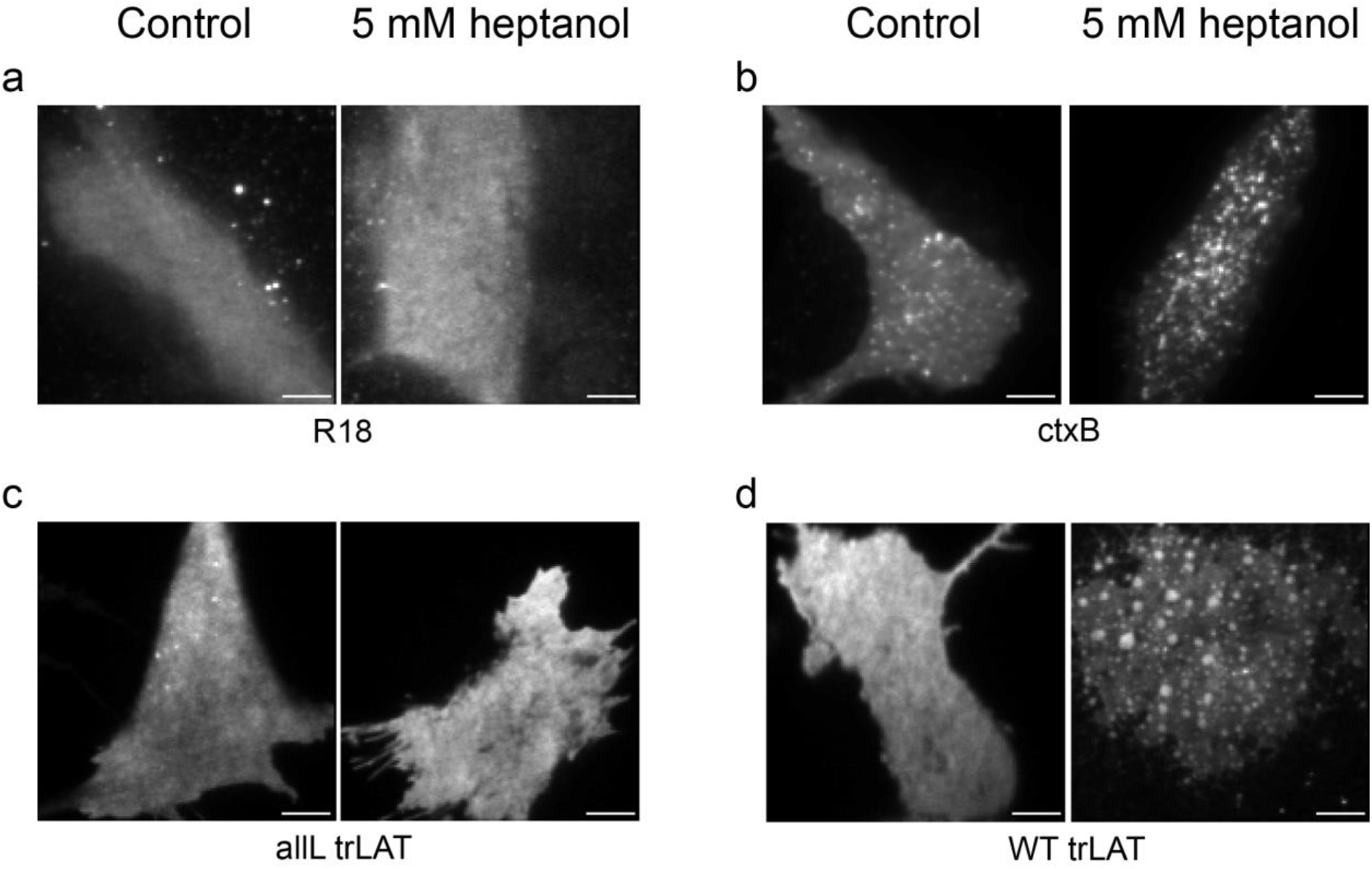
Heptanol phase separates molecules based on their domain preference. Representative TIRF images of probe organization on SH-SY5Y cells before and after 5 mM heptanol treatment. **(a)** Rhodamine 18 (R18). **(b)** Cholera toxin B (CTxB). **(c)** allL-trLAT. **(d)** WT-trLAT. Experiments were repeated at least three times. The scale bars represent 5 μm.

Our results show that the lateral segregation of proteins/markers that preferentially reside in the L_o_ phase is affected in response to treatment with heptanol, and these molecules form microscopic clusters in the cell membrane. This is in accordance with our hypothesis that hyperfluidization of the plasma membrane induced by heptanol leads to stabilized phase separation in membranes. These experiments show that heptanol mediated phase separation allows determining the phase preference of molecules that are transfected in cells or are labelled directly by an extrinsic label as in case of CTxB.

### Domain preference of signaling related proteins as determined by heptanol induced phase separation

After validating that heptanol treatment can identify the phase preference of marker proteins in live cell membranes, we used this method to determine the phase preference of signaling-related proteins. We specifically focused on those proteins for which pre-existing knowledge regarding their phase preference is available.

H-Ras and K-Ras are GTPases that function as molecular switches in the transduction of extracellular signals to the cytoplasm and nucleus. The membrane anchor in H-Ras consists of palmitoylated cysteine residues, thus directing it to the L_o_ phase in the membrane^50^. In contrast, K-Ras consists of a polybasic domain of six contiguous lysines, which confers L_d_ preference to this protein^50^. We transfected the SH-SY5Y cells with H-Ras and K-Ras constructs and treated the cells with heptanol. In untreated cells, H-Ras and K-Ras showed homogenous expression over the whole cell membrane. In heptanol-treated cells, as expected, H-Ras undergoes clustering upon heptanol treatment, while K-Ras does not (Figure 3a,b).

**Figure 3:**
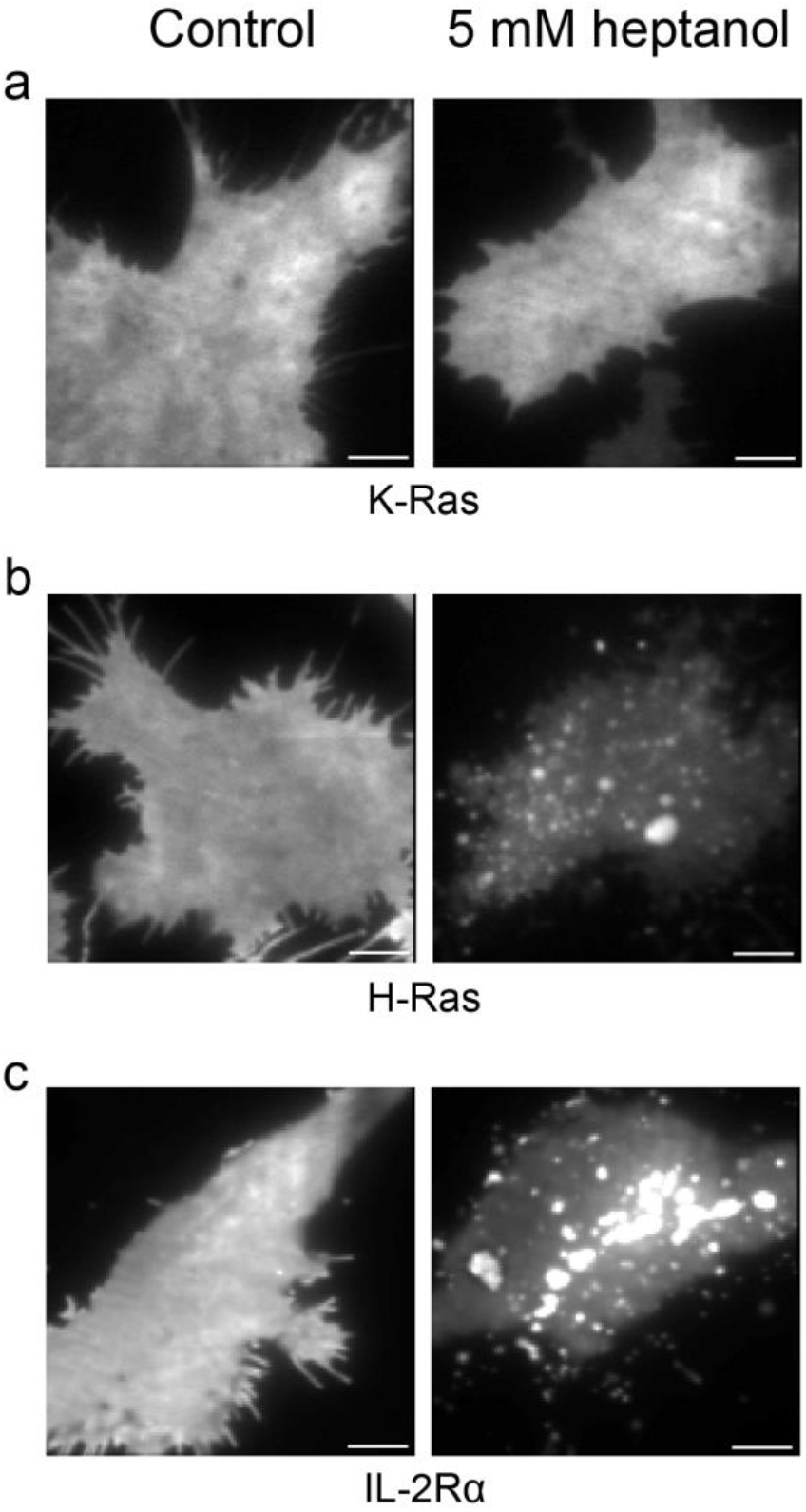
Domain preference of signaling related proteins as determined by heptanol-induced phase separation. Representative TIRF images of protein organization on SH-SY5Y cells before and after 5 mM heptanol treatment. **(a)** K-Ras. **(b)** H-Ras. **(c)** IL-2Rα. Experiments were repeated at least three times. The scale bars represent 5 μm.

Interleukin-2 (IL-2) signaling is directly regulated by the differential localization of IL-2Rβ-IL-2Rγ complexes in the soluble fraction of the membrane, i.e., L_d_ phase, while IL-2Rα is enriched in lipid microdomains, i.e., L_o_ phase of the membrane. Upon activation, three subunits of IL-2R heterotrimerize in the soluble fraction of the membrane^51^. Since phase preference of IL-2Rα is known, we subjected it to heptanol treatment and imaged changes in its lateral organization. IL-2Rα was homogenously expressed in untreated SH-SY5Y cells (Figure 3c). As expected, heptanol treatment-induced cluster formation also in this case (Figure 3c), thus validating our hypothesis.

Further, we probed the localization of epidermal growth factor receptor (EGFR), a well-studied protein; however, its membrane organization is not fully known. EGFR has been shown to localize in cholesterol-dependent domains^10,40,52–54^. However, previous work from our group showed that EGFR only partially depends on cholesterol-dependent domains^10^, thus indicating the presence of EGFR molecules also resides in the cholesterol independent fraction of the cell membrane. To gain more insights regarding the EGFR localization, we treated SH-SY5Y cells expressing EGFR with heptanol. Our results show that heptanol treatment leads to the formation of more clusters than those that were present before the treatment (Figures 4a,b, S1,S2a,b).

**Figure 4:**
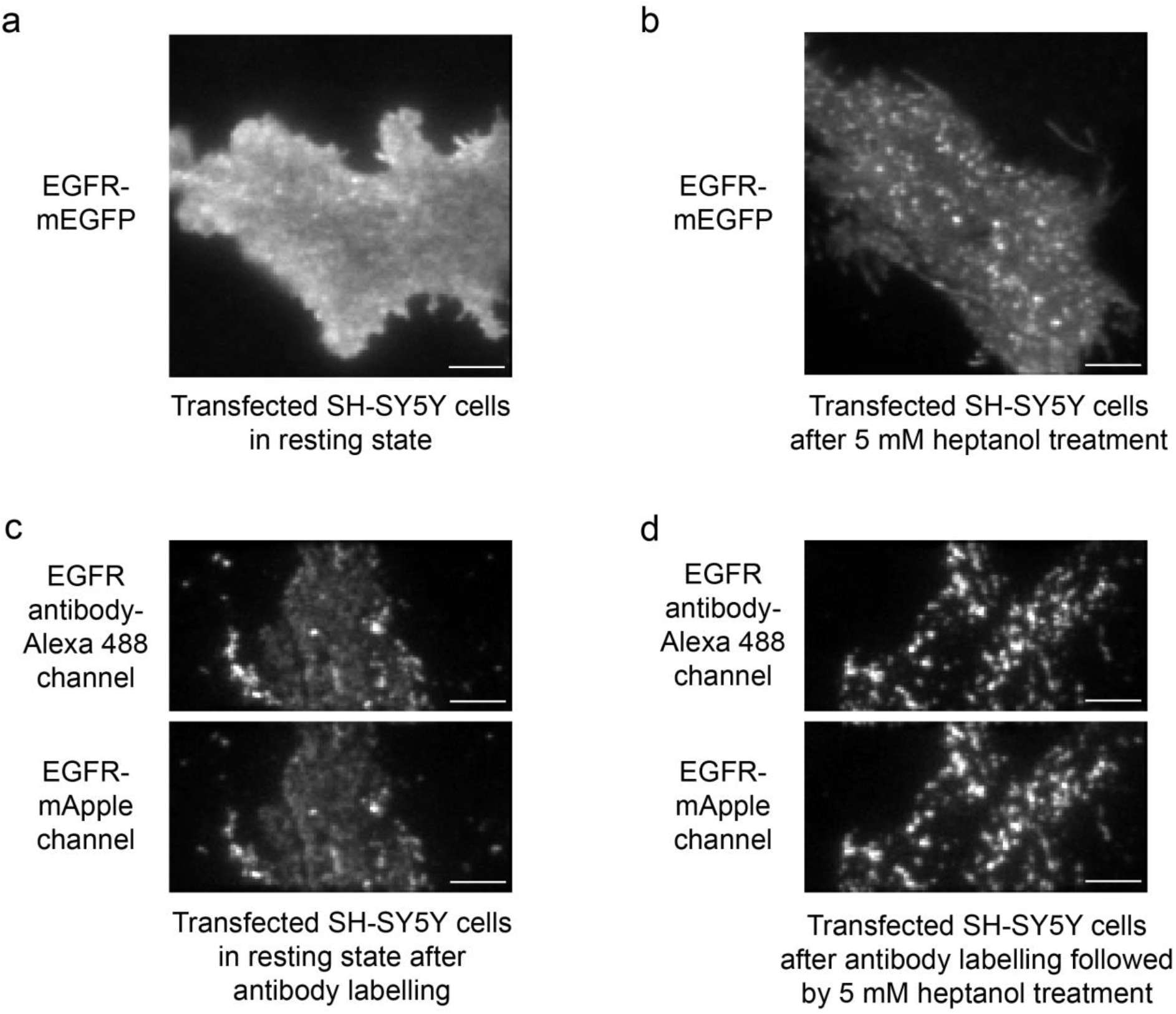
Heptanol-induced phase separation of EGFR. Representative TIRF images of EGFR transfected SH-SY5Y cells under various conditions. **(a)** EGFR-mEGFP in resting transfected cells. **(b)** EGFR-mEGFP after 5 mM heptanol treatment. **(c)** EGFR antibody-Alexa 488 and EGFR-mApple channels after labelling of resting cells with EGFR antibody. **(d)** EGFR antibody-Alexa 488 and EGFR-mApple channels after labelling of the cells with EGFR antibody followed by 5 mM heptanol treatment. Experiments were repeated at least three times. The scale bars represent 5 μm.

### Comparison of cluster area fraction and cluster sizes for various probes

We compared the area fraction covered by clusters and cluster sizes on cell membranes expressing various probes – GPI, WT-trLAT and EGFR – before and after heptanol treatment (Figure S1). In resting cells, the samples exhibit almost homogeneous lateral segregation of molecules and clusters cover 3±3%, 2±0.1% and 2±2% of the total cell membrane area for GPI, WT-trLAT and EGFR, respectively. After heptanol treatment, the area fraction covered by clusters increased to 20±8%, 37±3% and 10±2%, respectively (p<0.5 in all cases; refer Figure S1). The smaller increase for EGFR compared to GPI and WT-trLAT could be because of a population of EGFR that resides in the L_d_ phase^10,40,53^. However, this requires further investigation.

All the samples showed a similar range of 0.6-1 μm cluster size in both resting and treated cells (p>0.5; refer Figure S1). It is important to note that since the cluster area fraction is <5% in resting cells, which is significantly less than the cluster area fraction measured post heptanol treatment, these values correspond to the size of some rarely occurring clusters on the membrane.

EGFR was also labelled with an antibody conjugated with Alexa 488 dye. The antibody labelling induced some clustering (12±1%) of EGFR (Figures 4c, S1, S2c). The clustering increased upon heptanol addition (24±5%; p < 0.5) indicating heptanol-induced phase separation (Figures 4d, S1, S2d). Therefore, MFIC can also be used with extrinsic fluorescent tags and does not require genetic labeling.

### Heptanol based assay to determine phase preference of membrane molecules is fast and reversible

Although microscopic clustering of molecules is typically observed 10-15 minutes after the treatment, the immiscibility of phases starts immediately after the addition of heptanol. This was revealed by our ITIR-FCS measurements consisting of 300,000 frames with an exposure time of 1 ms, i.e., dynamics readouts of 5 minutes. In these FCS videos^55^, the first 50,000 frames were recorded on untreated cells, and then cells were treated with heptanol while recording the data. The heptanol-induced changes in membrane dynamics were then recorded for the subsequent 250,000 frames. We performed these experiments on WT-trLAT and trLAT-allL expressing SH-SY5Y cells and then utilized diffusion maps calculated from consecutive, non-overlapping sections of 50,000 frames.

In the case of WT-trLAT, the first 50,000 frames show a homogeneous diffusion in the cell membrane. Upon heptanol treatment, there was an increase in D with time, indicating an increase in fluidity (Figure S3a). Moreover, we observed that from 200,000 frames onwards, pixels appeared with high D, indicated by the separation of the high D pixels (green and red) from the cluster of pixels with low D (blue). However, TIRF images constructed from the projection of the last 50,000 frames did not show any noticeable microscopic clustering. Clustering of molecules was observed 10-15 minutes post heptanol treatment, only. This implies that smaller clusters, which are difficult to observe via imaging, are detected by the effect on molecular dynamics already early on.

Similar to WT-trLAT, trLAT-allL showed homogeneous diffusion in the first 50,000 frames (Figure S3b). And heptanol treatment increased the overall D of the molecules. But unlike WT-trLAT, in this case, there was no noticeable phase separation as even after 250,000 frames diffusion remained homogenous, indicating the absence of phase separation on the cell membrane.

Subsequently, we tested if the clustering of molecules induced by heptanol-mediated hyperfluidization is reversible. For this purpose, we treated the cells with heptanol for 15 minutes, sufficient for cluster formation to occur. Then we washed the heptanol solution from the cell membranes and incubated the cells for 24 hours in complete growth medium at 37 °C with 5% CO_2_. We observed that 24 hours after washing away the heptanol solution, the clusters had disappeared almost completely (Figure S4), demonstrating the reversible nature of this assay. However, it is to be noted that currently we do not know the full nature of the clusters and whether protein interactions are involved in addition to phase separation in their formation.

Furthermore, we asked if this result is obtainable in cell membranes other than SH-SY5Y. Hence, we transfected WT-trLAT and trLAT-allL in Hela cells and examined their lateral organization before and after heptanol treatment. As observed in SH-SY5Y cell membranes, WT-trLAT undergoes clustering upon heptanol treatment (Figure 5a) while trLAT-allL does not show any microscopic changes in their organization in Hela cell membranes (Figure 5b).

**Figure 5:**
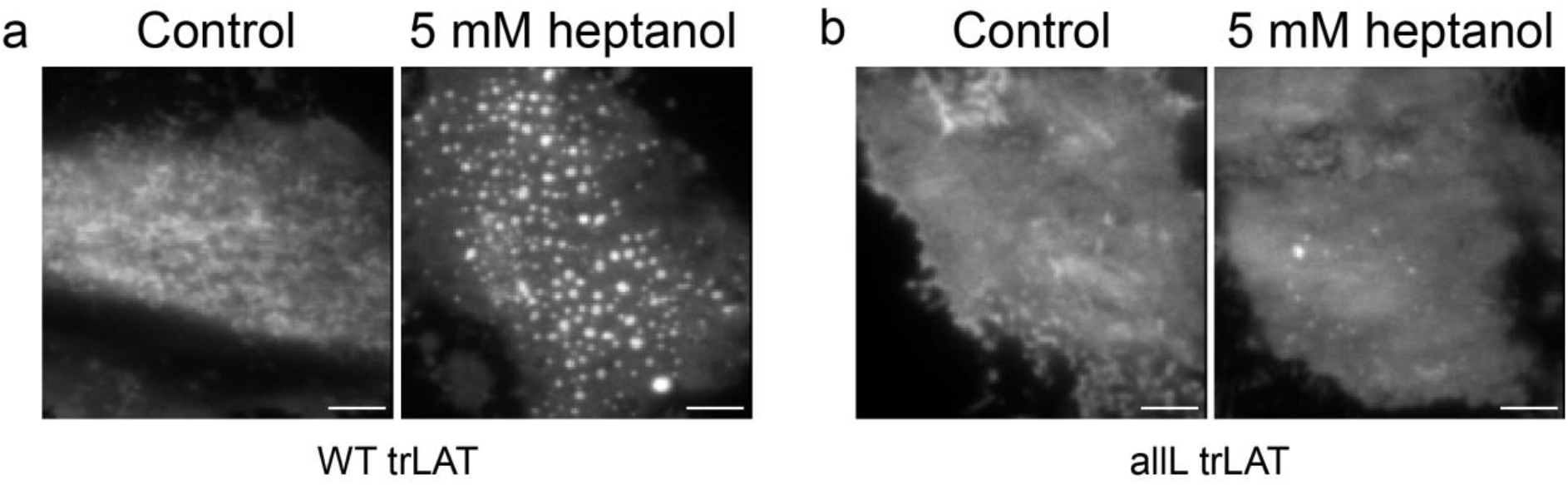
Validation of heptanol-based method for phase preference determination in HeLa cells. Representative TIRF images of protein organization on HeLa cells before and after 5 mM heptanol treatment. **(a)** WT-trLAT. **(b)** all-trLAT. Experiments were repeated at least three times. The scale bars represent 5 μm.

Most routinely used methods to determine membrane localization of molecules such as detergent-resistant membrane extraction, membrane extraction followed by mass spectrometry, and immunostaining are time consuming and are artifact-prone. Using membrane-fluidizing agents allows the determination of membrane localization of molecules in a few minutes and in a live-cell membrane environment, thus providing a more physiologically relevant result. This method can be implemented on genetically fluorescence labeled as well as on endogenously expressed molecules labelled extrinsically as demonstrated by experiments performed on CTxB and EGFR. This method will aid in the quick identification of protein phase preference in cell membranes. Moreover, heptanol-mediated phase separation in cell membranes can be utilized to probe the thermodynamic properties of cell membranes by computational methods as investigated for other small molecules^56^.

## CONCLUSION

Here, we have developed and validated a novel live-cell assay, membrane fluidizer-induced clustering (MFIC), for determining the phase localization of proteins in cell membranes using heptanol. We demonstrated that heptanol induces membrane fluidization and stabilizes phase separation in cell membranes resulting in formation of microscopic clusters of L_o_ domains which are easy to detect. To test the ability of this assay in determining the molecule phase preference, we carried out heptanol treatment on fluorescently labelled lipids and proteins for which phase preference is known. Compared to the existing methods of ascertaining molecule phase preference, this method is fast as one can get the result in 15-20 minutes post heptanol treatment. Moreover, this method is less artifact-prone as it is carried out in live cell membranes.

## ACKNOWLEDGEMENTS

T.W. gratefully acknowledges funding from the Singapore Ministry of Education (R-154-000-B53-114). D.L. is the recipient of a scholarship (No. 201906145023) from the China Scholarship Council. H.B. is the recipient of a research scholarship of the National University of Singapore.

## AUTHOR CONTRIBUTIONS

T.W. and A.G. conceived and designed the study. T.W. supervised the study and provided the resources. A.G., D.L., H.B. and Z.C. performed the experiments. A.G., D.L. and H.B. analyzed the data. A.G., H.B. and T.W. wrote the manuscript.

## SUPPLEMENTARY FIGURES

**Figure S1:**
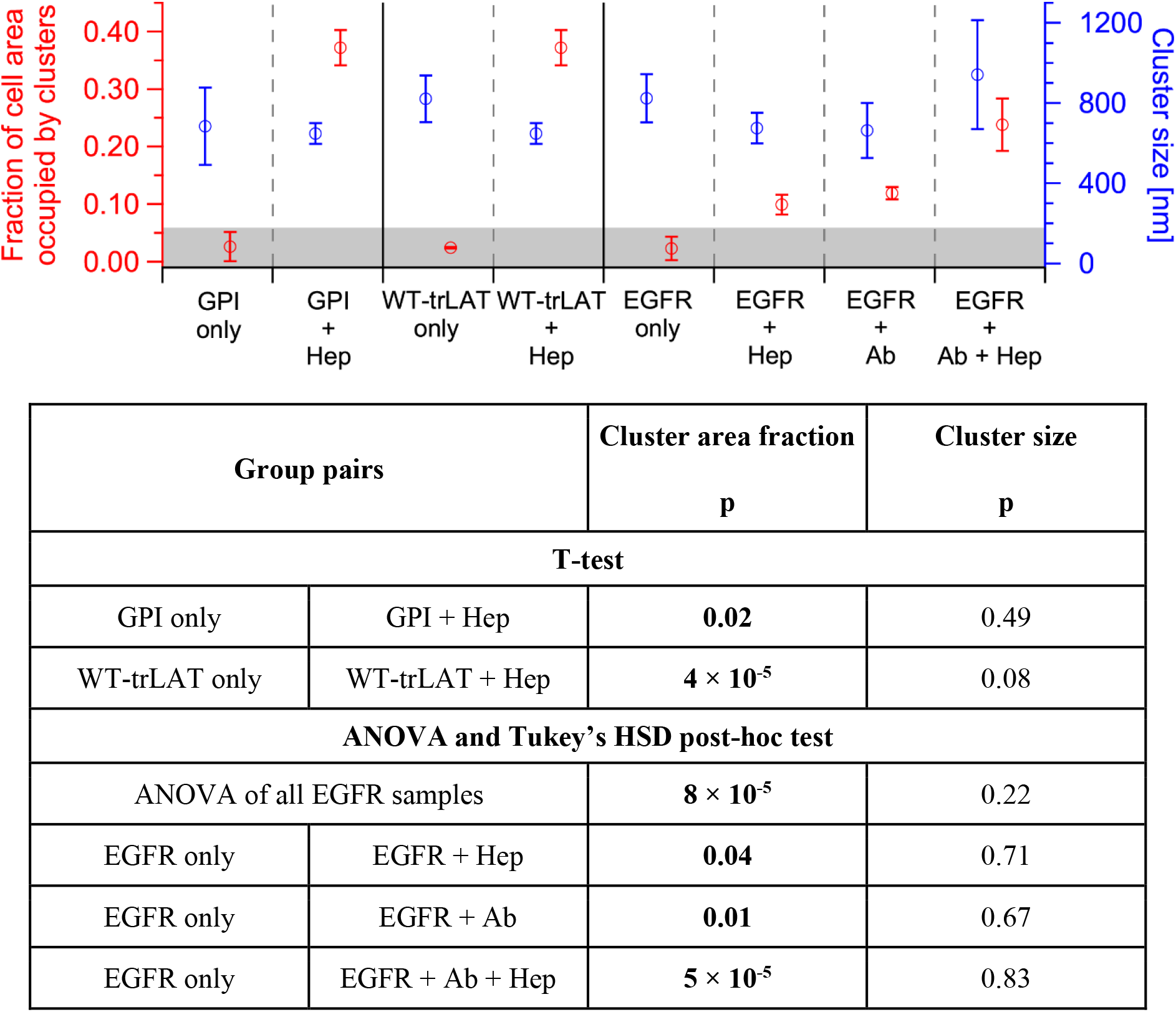
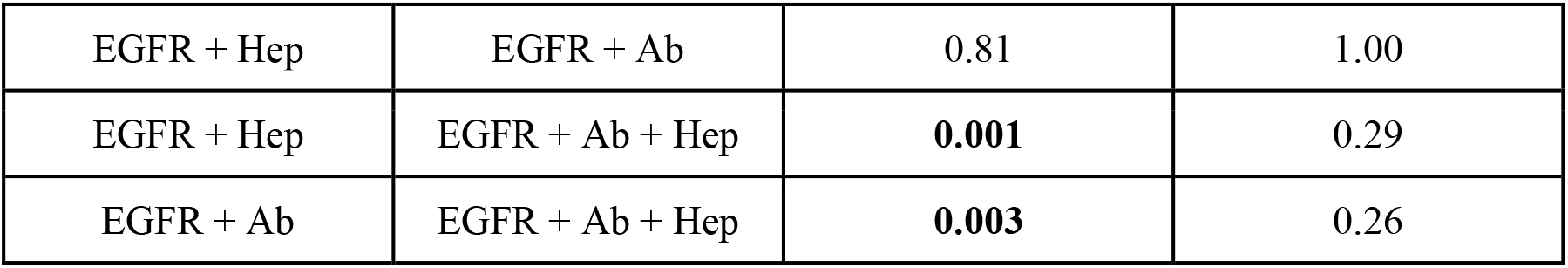
Comparison of fractional area covered by clusters on cell membranes for various probes. Cluster area fractions and cluster sizes for GPI, WT-trLAT, and EGFR after various treatments (Hep: heptanol, Ab: antibody) are shown here. Before heptanol treatment the membranes are homogeneous with negligible clustering (<5% - grey area). However, cluster sizes measured before heptanol treatment are comparable with the ones measured after the treatment. The statistical analyses (refer Materials and Methods ➔ Data analysis) were performed on three images recorded from three independent experiments. Mean ± SD is shown here. The significant differences (p < 0.05) are highlighted in bold.

**Figure S2:**
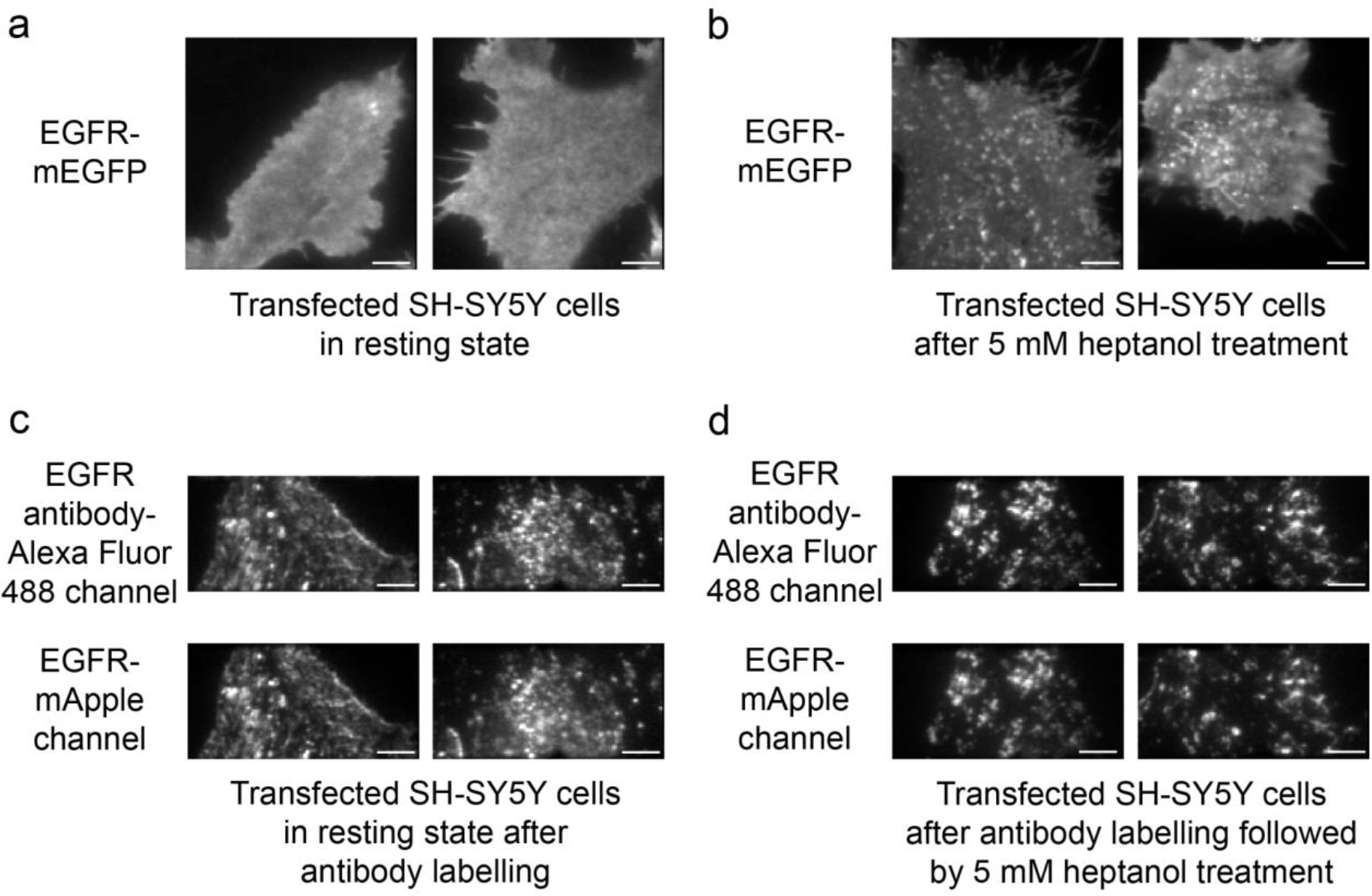
Replicates of heptanol-induced phase separation of EGFR. Representative TIRF images of transfected EGFR in SH-SY5Y cells from replicates of experiments shown in Figure 1. **(a)** EGFR-mEGFP in resting transfected cells. **(b)** EGFR-mEGFP after 5 mM heptanol treatment. **(c)** EGFR antibody-Alexa 488 and EGFR-mApple channels after labelling of resting cells with EGFR antibody. **(d)** EGFR antibody-Alexa 488 and EGFR-mApple channels after labelling of the cells with EGFR antibody followed by 5 mM heptanol treatment. Experiments were repeated at least three times. The scale bars represent 5 μm.

**Figure S3:**
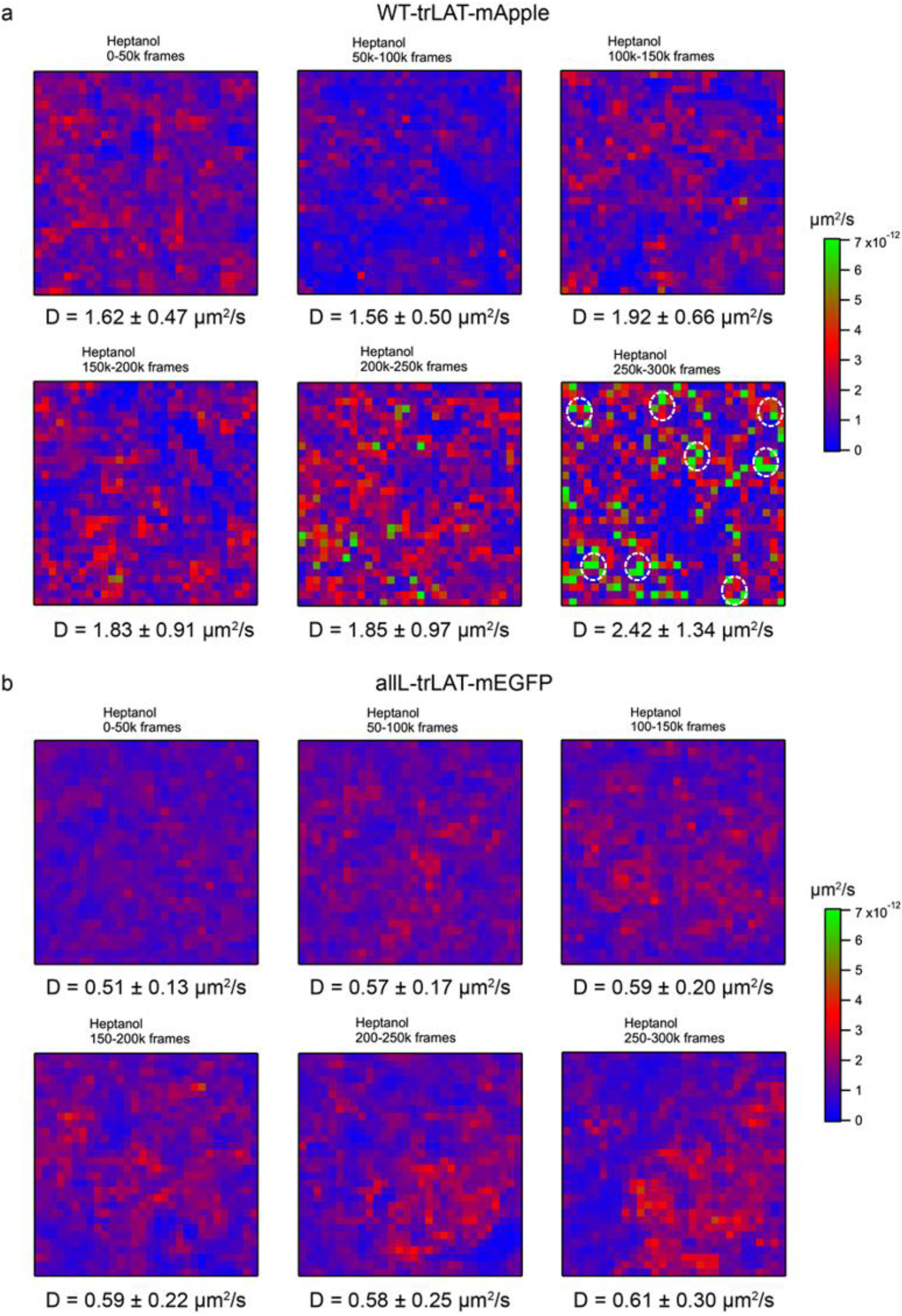
Diffusion changes after heptanol treatment monitored by ITIR-FCS. Representative diffusion maps obtained by ITIR-FCS measurements performed on SH-SY5Y cells expressing **(a)** WT-trLAT-mApple and **(b)** allL-trLAT-mEGFP. Measurements consist of 300,000 frames with an exposure time of 1 ms. 5mM Heptanol was added to the cells at 50,000 frames. The diffusion maps are calculated from non-overlapping 50,000 frames. Average diffusion coefficient (D) corresponding to each diffusion map are noted underneath the maps. The dotted white circles on the diffusion map obtained from correlating 250,000-300,000 frames in WT-trLAT-mApple expressing cells (last panel in (a)) mark the group of pixels exhibiting higher D clustered together in the cell membrane.

**Figure S4:**
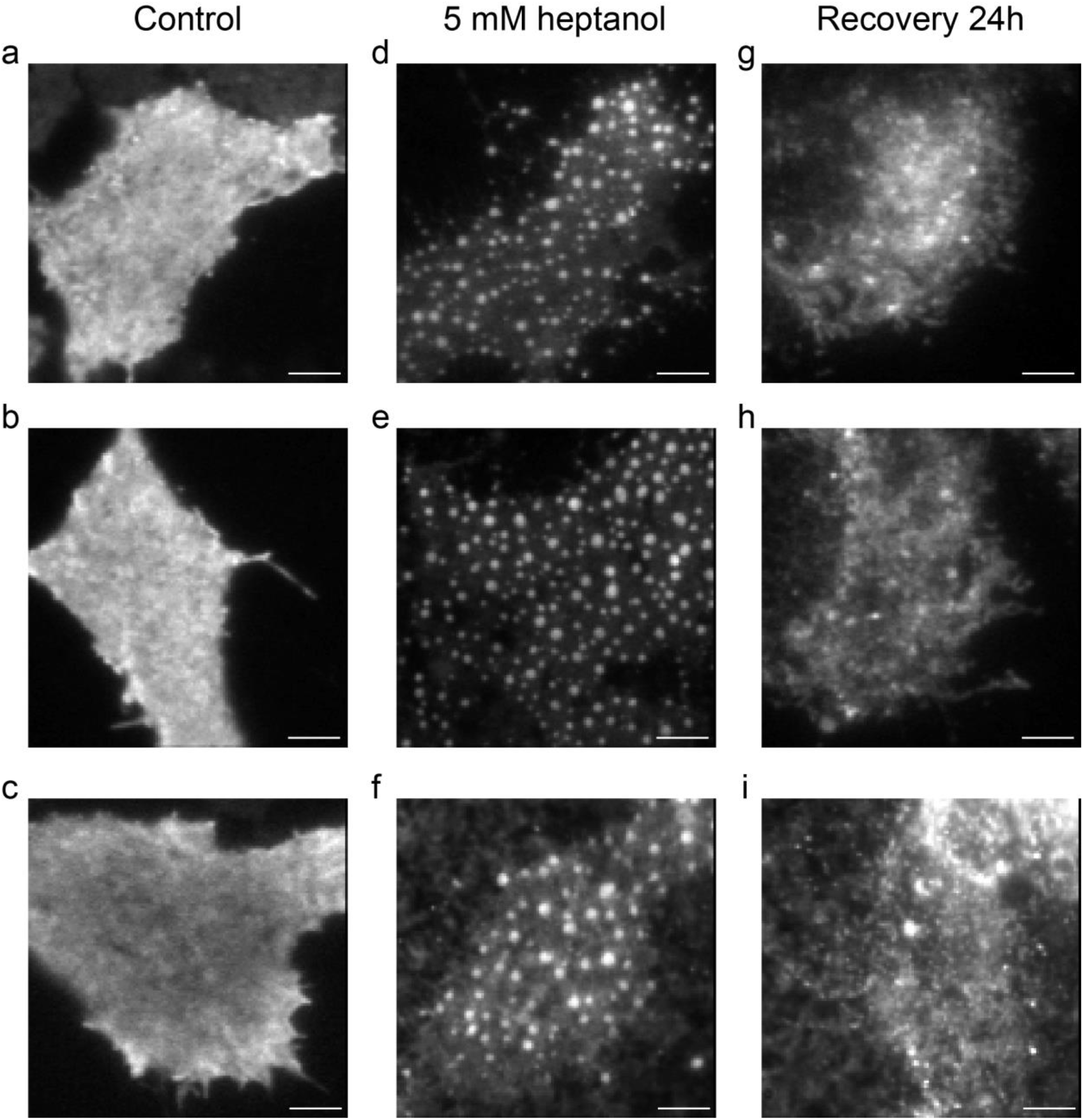
Heptanol-induced domain reorganization is reversible. Representative TIRF images of SH-SY5Y cells expressing GFP-GPI under various conditions. (a,b,c) Resting transfected cells. (d,e,f) Cells treated with 5 mM heptanol for 15 minutes. (g,h,i) Cells 24 hours after washing away the heptanol solution and replacing it with complete growth medium. These images are from three independent experiments. The scale bars represent 5 μm.

